# *Drosophila* macrophage self-renewal is regulated by transient expression of PDGF- and VEGF-related factor 2

**DOI:** 10.1101/2020.08.24.255638

**Authors:** Daniel Bakopoulos, James C. Whisstock, Coral G. Warr, Travis K. Johnson

## Abstract

Macrophages are an ancient animal blood cell lineage critical for tissue homeostasis and defence against pathogens. Until recently, their numbers were thought to be sustained solely by specialised hematopoietic organs. It is now clear that many macrophages are instead replenished by self-renewal, yet the signals that regulate this remain poorly understood. In *Drosophila melanogaster*, macrophages (known as plasmatocytes) undergo a phase of rapid population expansion via self-renewal, making *Drosophila* an attractive model for revealing the signals and regulatory mechanisms involved. However, no central self-renewal pathway has been identified in *Drosophila*. Here, we investigated the PDGF-/VEGF-receptor pathway as a candidate for playing this role. Analysis of larvae deficient for each of the three PDGF-/VEGF-receptor ligands Pvf1-3 revealed Pvf2 as a major driver of macrophage self-renewal in *Drosophila*. We further found that only a small proportion of blood cells express *Pvf2*, and knockdown experiments implicate these cells as a major source of *Pvf2* in self-renewal. Lineage tracing studies support the idea that *Pvf2* expression in blood cells occurs transiently throughout the macrophage self-renewal period, and in response to an as yet unidentified cue. These data define the regulation of *Pvf2* expression in blood cells as a central mechanism by which macrophage self-renewal is controlled. Given the strong parallels that exist between *Drosophila* and vertebrate macrophage systems, it is likely that similar mechanisms are at play across animal phyla.

## Introduction

Macrophages are highly versatile blood cells that reside within body tissues and play critical roles in immune surveillance, pathogen and cellular debris clearance, inflammation, wound healing, and a variety of developmental processes (*1-3*). Macrophage numbers are highly dynamic during the lifetime of an animal. For example, their numbers increase upon infection as part of the immune response, while during development their populations expand rapidly in order to colonise growing tissues for important local homeostatic duties (*1, 4-6*). For many years, the vertebrate macrophage population was thought to be sustained solely by the contributions of blood cell progenitors and hematopoietic stem cells (HSCs). Recently however, it has become clear that the ability of macrophages to proliferate (or self-renew) locally within tissues also plays a major role (*1, 7*). Despite the importance of self-renewal to the macrophage population, the signals and pathways that control this process remain poorly understood (*8, 9*). Adding to the complexity is that often the vertebrate self-renewing macrophage populations coexist with those of HSC origins, thereby making the task of identifying novel self-renewal mechanisms challenging (*10, 11*).

The blood cell systems of many invertebrates, including *Drosophila melanogaster*, similarly comprise macrophages (called plasmatocytes) that arise de novo from blood cell progenitors or via self-renewal (*12-14*). In *Drosophila*, macrophages make up more than 90% of the total population of blood cells (known as hemocytes) (*6*). The remainder are crystal cells (<5%), responsible for protective melanisation reactions during infection, and lamellocytes, which appear upon wasp parasitisation to encapsulate wasp eggs (*6, 15-17*).

All *Drosophila* blood cells originate from two populations of blood cell progenitors (*18, 19*). During embryogenesis, the first blood cell progenitor population gives rise to both crystal cells and macrophages (*12, 20*). The latter persist into the larval stages where they can either circulate throughout the blood (called lymph) or reside in tissues termed hematopoietic pockets (*19-21*). During the subsequent four days that constitute the larval stage, the macrophage population expands dramatically from several hundred cells to approximately 10,000 via self-renewal (*6, 13, 19, 22*). During this time, crystal cells and lamellocytes arise from the macrophage pool via transdifferentiation (*5, 23*).

The second population of blood cell progenitors originate within a HSC-like niche in the larva called the lymph gland (*14*). The blood cells that are produced here, however, are not released into circulation until the onset of metamorphosis, which marks the completion of the larval stage (*19, 24*). This means that, unlike in vertebrates, the self-renewing macrophages in fly larvae are easily separated from those of HSC origin. For this reason, *Drosophila* is emerging as a useful model for the study of self-renewal mechanisms (*11, 25*). Despite this, a central cell signalling pathway responsible for driving macrophage self-renewal in remains *Drosophila* to be described. Thus, our understanding of the mechanism(s) by which this process is controlled remain incomplete.

A candidate for this is the PDGF*-* and VEGF*-*receptor related (Pvr) pathway. Pvr is a receptor tyrosine kinase with three known ligands, PDGF*-* and VEGF*-* related factors 1-3 (Pvfs1-3), and is the sole *Drosophila* orthologue of the PDGF and VEGF family of receptors (*26, 27*). Pvr signalling has been previously implicated in the survival of blood cells during embryogenesis, and while it is thought to also influence larval blood cells, its role here has remained unclear (*28-31*).We therefore investigated the Pvr pathway and found it to be a major driver of macrophage self-renewal in *Drosophila*. We find that final larval macrophage numbers are drastically reduced in *Pvf2* and *Pvf3* mutants and show that *Pvf2* alone influences macrophage self-renewal. We further find that *Pvf2* is transiently expressed in only a small proportion of blood cells, and that its modulation here causes large and concomitant changes in their population size. These data support a new model for larval macrophage self-renewal in *Drosophila* that involves the transient transcriptional regulation of *Pvf2* in blood cells.

## Materials and methods

### *Drosophila* stocks and maintenance

The following stocks from the Bloomington *Drosophila* Stock Centre were used: *w*^1118^ (BL3605), *hml-GAL4,UAS-GFP* (BL6397), *hmlΔ-GAL4,UAS-GFP* (BL30142), *Pvf1*^*1624*^ (BL11450), *Pvf2*^*MI00770*^ (BL32696), *Pvf3*^*MI04168*^ (BL37270; called *Pvf3*^*MiMIC*^ here), *df*^*BSC291*^ (BL23676), *UAS-Pvf2* (BL19631), *UAS-eYFP;Sp/CyO;Dr/TM3* (BL60291), *hs-Cre,vas-dϕC31* (BL60299), *Sp/CyO;lox(Trojan-GAL4)x3* (BL60311), *SrpHemo-H2A-mCherry/CyO;Dr/TM3* (BL78361), *UAS-EOS-GFP* (BL32228), *UAS-FLP,UAS-RFP,Ubi*^*p63*^*-FRT-STOP-FRT-GFP* (BL28280, called *G-TRACE* here); *UAS-RFP* (BL8546). We obtained *UAS-Pvf2 RNAi* (13780R-1) from NIG Japan, and *UAS-eYFP* was derived from BL60291. T2A-GAL4 lines were generated via RMCE using BL60291, BL60299 and BL60311 as previously described (*32*). *Pvf2*^*T2A-GAL4*^, *Pvf3*^*MiMIC*^, *df*^*BSC291*^, *UAS-Pvf2* and *UAS-Pvf2 RNAi* were maintained over green (GFP) balancers. *hmlΔ-GAL4,UAS-GFP* was maintained over TM6B. *hmlΔdsRed* was a gift from Katja Brückner (*13*). Flies were raised and maintained on standard sucrose, yeast and semolina fly media. All experiments were performed at 25 □C.

### Quantitative RT-PCR

RNA was extracted from the anterior half of wandering larvae using Trisure reagent (Bioline) and cDNA synthesised using oligodT and random hexamers (Bioline). Quantitative PCR was performed on a lightcycler (Roche) with SYBR chemistry. The following primers pairs were designed (PerlPrimer) (*33*), validated and tested for efficiency: *Pvf2:* F 5’-TGA AAG AGC GAA TCG CCG AAC AA-3’ and R 5’-GCA GAT ACC CTC CTT TGC CAT CA-3’, *Pvf3*: F 5’-TCT ATA CGC CTC ACT GCA CCA TCC-3’ and R 5’-ACT GCG ATG CTT ACT GCT CTT CAC-3’. Reactions for *Pvf2* and *Pvf3* and the control gene *cyclin K:* F-5’-GAG CAT CCT TAC ACC TTT CTC CT-3’, R-5’-TAA TCT CCG GCT CCC ACT G-3’ (*34*), were run in triplicate for 3-4 biological replicates per genotype and fold changes determined using the ΔΔCT method. Differences between means were assessed using an unpaired t-test with Welsch’s correction (Graphpad Prism 8).

### Larva rearing and weighing

For wandering third instar larvae, 0-24 hour old larvae were collected from egg-lays on apple juice agar supplemented with yeast paste, then reared on standard fly media under non-crowded conditions until the wandering stage. Where newly moulted third instar larvae were needed, developing larval cultures were suspended with sucrose (20% w/vol) and second instar larvae placed onto fresh fly media based on their anterior spiracle morphology. This was repeated two hours later and the newly moulted third instar larvae used in experiments. For first instar larvae, 2 hour egg-lays were performed and individuals allowed to develop a further 44-46 hours on apple juice agar supplemented with yeast paste before bleeding. Correct genotypes were selected using balancers marked by GFP expression or Tb as required. To weigh larvae individual wandering third instar larvae were washed in phosphate buffered saline (PBS), checked for debris, blotted dry and weighed on an ultra-microbalance (XP2U, Mettler Toledo) immediately prior to bleeding.

### Blood cell quantification

Blood cells were extracted from third instar wandering stage larvae (unless otherwise stated) carrying a fluorescent blood cell marker – *hml-GAL4,UAS-GFP* (*35*), *hmlΔ-GAL4,UAS-GFP* (*36*) or *hmlΔdsRed* (*13*) – or *G-TRACE* transgenes as previously described (*22*). Briefly, circulating blood cells were bled from individual larvae in PBS in an 8-well slide (Ibidi) for at least two minutes through from a hole torn in the dorsal-posterior cuticle. Tissue-resident blood cells were removed by scraping the remaining blood cells from the carcass into a new well, while taking care not to disrupt the lymph gland. Cells were resuspended by pipetting to minimise clumping and allowed to settle for at least 10 minutes. The entire well was imaged using an Olympus CV1000 for all experiments, except for the *Pvf2* knockdown experiment where a Leica AF6000 LX was used. The threshold feature on ImageJ was used to determine the number of relevant classes of fluorescent cells per well, applying the same threshold across all images in a given experiment (*37*). Differences between means were assessed using the non-parametric Mann-Whitney test (Graphpad Prism 8). When larval weights or stage differences were accounted for, regression analyses were performed following log transformation to ensure linearity across the data range (RStudio) as previously described (*38*).

Crystal cells were counted using the black cell assay (*39*). Briefly, wandering third instar larvae were placed into 100μL H_2_O and heated in a 65°C water bath for 20 minutes. Larvae were then stored in H_2_O at -20°C for no longer than 72 hours before being imaged on a Leica M165 FC. The number of crystal cells on the dorsal side of the four posterior-most abdominal segments were counted in ImageJ using the Cell Counter plugin for ≥13 larvae per genotype (*37*). Differences between means were assessed using the non-parametric Mann-Whitney test (Graphpad Prism 8).

### Blood cell self-renewal and death measurements

Self-renewal rates were assayed by feeding newly moulted third instar larvae carrying *hmlΔdsRed* standard fly media containing 200μM 5-ethynyl-2’-deoxyuridine (EdU), 1:50 dimethyl sulfoxide (DMSO) and red food colouring for 4 hours prior to bleeding, as previously described (*40*). Larvae that had ingested food were selected and their blood cells extracted into a droplet of PBS on a Conconavalin A (ConA, 0.5mg/mL) coated coverslip and allowed to settle for 30 minutes. Blood cells were fixed and stained using the Click-iT EdU Proliferation Kit for Imaging (Invitrogen), imaged and quantified as described above. For cell death measurements, cells were bled from newly moulted third instar larvae and adhered to ConA coverslips. Bled cells were fixed and stained using TUNEL reagents (Roche) before imaging and quantification. Differences between means were assessed using the non-parametric Mann-Whitney test (Graphpad Prism 8).

### Embryonic blood cell imaging

Embryos were dechorionated for 3 minutes in 50% (vol/vol) bleach and fixed in a mixture of 1:1 PBS containing 4% paraformaldehyde (PFA) and heptane for 30 minutes before methanol devitellinisation. Embryos were rehydrated in PBS containing 0.1% Triton X (PBS-T), blocked for 30 minutes in PBS-T with 5% (vol/vol) normal goat serum (G-9023, Sigma) before overnight incubation with anti-GFP (1:1000, A-6455 Invitrogen). Anti-rabbit secondary (1:500, A-11034 Invitrogen) was applied and embryos mounted in Vectashield before imaging. For live-imaging, embryos were dechorionated for 90 seconds and mounted in 8-well slides (Ibidi) under PBS for imaging.

### Lineage tracing with EOS-GFP and live imaging of larvae

Third instar larvae expressing EOS-GFP were irradiated in H_2_O using a DAPI filter and 5x objective (Leica DMLB compound) for at least 2 minutes to ensure high conversion efficiency while maximising larval survival. Larvae were then placed on standard fly media. Larvae were imaged live 24 hours later, on a microscope slide held in place by double-sided tape under a coverslip (towards the dorsal side) as previously described (*23*).

## Results

### Larval macrophage number is influenced by *Pvf2* and *Pvf3*, but not *Pvf1*

To determine whether any of the three known Pvr ligands (Pvf1-3) may play a central role in the control of macrophage population expansion in larvae, we sought to determine whether larvae mutant for each individual *Pvf* exhibit defects in macrophage number. For *Pvf1*, a validated null allele (*Pvf1*^*1624*^) was available (*27*), however this was not the case for *Pvf2* or *Pvf3*. We therefore generated null mutants for each of these genes by making use of available lines carrying MiMIC transposons inserted within their introns (Figure 1A). The MiMIC transposon arrests transcription when inserted in the correct orientation (*41*) and can be replaced with a variety of available cassettes by recombinase-mediated cassette exchange (RMCE) (*32*). For *Pvf2* the insertion (*Pvf2*^*MI00770*^) was incorrectly orientated, so we replaced the MiMIC transposon with a T2A-GAL4 cassette, which is intended to halt *Pvf2* transcription and instead produce GAL4 (*Pvf2*^*T2A-GAL4*^). The transposon located in *Pvf3* (*Pvf3*^*MI04168*^, called *Pvf3*^*MiMIC*^ here) was in the correct orientation and therefore expected to disrupt the gene. However, for the added utility of having a GAL4 reporter, we also replaced this with the T2A-GAL4 cassette (*Pvf3*^*T2A-GAL4*^). These insertions (MiMIC and T2A-GAL4) affect all known isoforms of *Pvf2* and *Pvf3* and likely truncate residual gene products to short fragments lacking their predicted receptor-binding PDGF domains (Figure 1A). Quantitative RT-PCR for the presence of exons downstream of the insertion sites in mutant larvae carrying the *Pvf2*^*T2A-GAL4*^ and *Pvf3*^*MiMIC*^ alleles confirmed that full-length transcripts are not detectable in the mutant lines (Figure 1B, C), indicating that these are likely null alleles.

**Figure 1.**
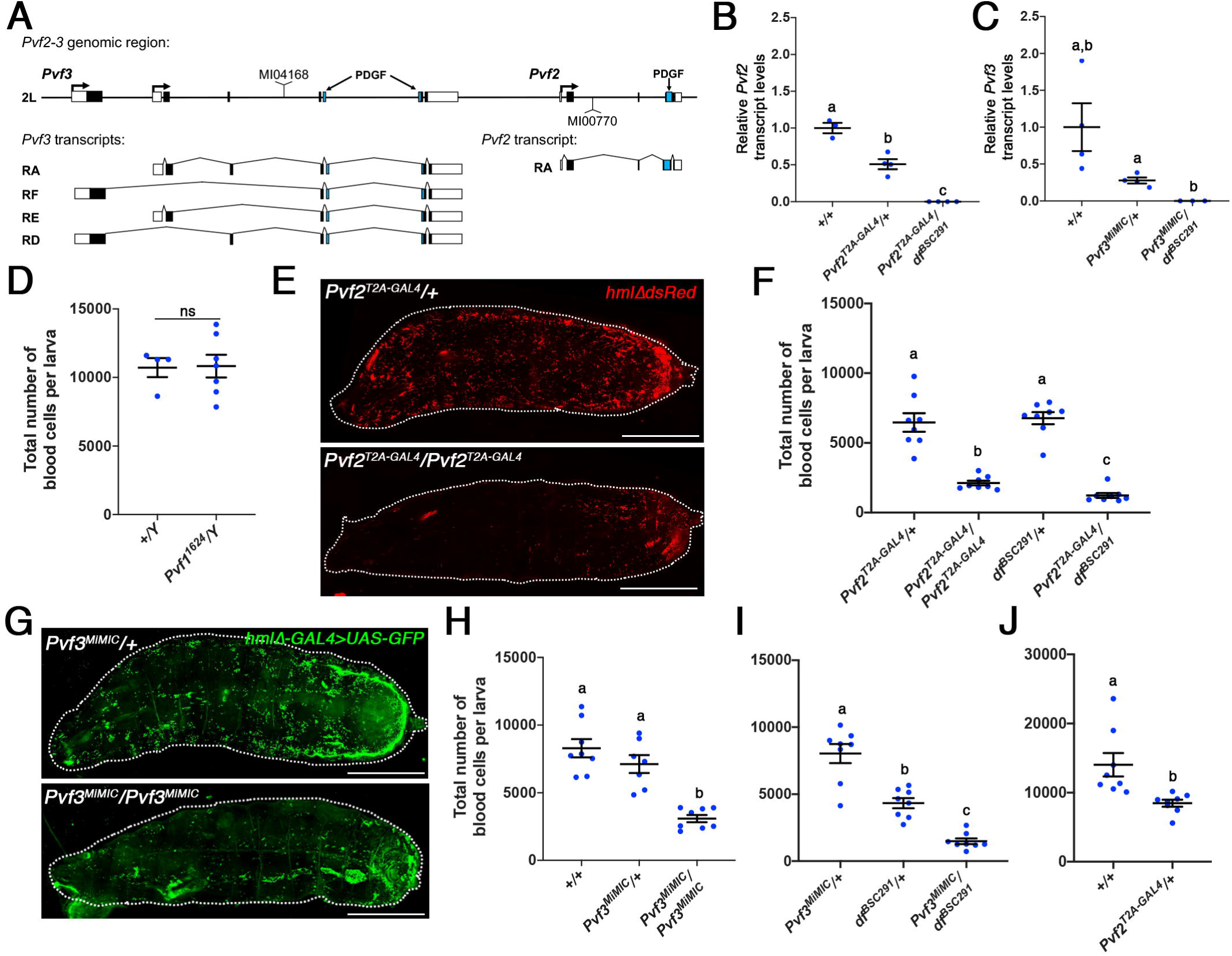
*Pvf2* and *Pvf3*, but not *Pvf1* influence larval blood cell number. (**A**) *Pvf2*-*3* genomic region and transcripts. Transcriptional start sites are indicated by arrows, boxes denote exons (coding sequence is filled with blue denoting PDGF domain sequences), and the positions of two relevant MiMIC transposons are shown. Relative transcript levels of *Pvf2* (**B**) and *Pvf3* (**C**) in larvae of the genotypes indicated shows that *Pvf2*^*T2A-GAL4*^ the *Pvf3*^*MiMIC*^ alleles fail to produce detectable transcript levels of *Pvf2* and *Pvf3*, respectively. (**D**) *Pvf1*^*1624*^*/Y* larvae exhibited no significant difference in their total blood cell number compared to wild-type controls (p>0.999, *hmlΔdsRed*, n≥4). (**E**) *Pvf2*^*T2A-GAL4*^*/Pvf2*^*T2A-GAL4*^ larvae appear to have fewer blood cells (*hmlΔdsRed*) than *Pvf2*^*T2A-GAL4*^*/+* controls. (**F**) Blood cells numbers in homozygous *Pvf2*^*T2A-GAL4*^ and *Pvf2*^*T2A-GAL4*^*/df*^*BSC291*^ larvae are strongly reduced compared to heterozygous controls (*hmlΔdsRed*, n=8). (**G**) *Pvf3*^*MiMIC*^*/Pvf3*^*MiMIC*^ larvae appear to have fewer blood cells (*hmlΔ-GAL4>UAS-GFP*) than *Pvf3*^*MiMIC*^*/+* controls. This was confirmed by blood cell quantification of *Pvf3*^*MiMIC*^*/Pvf3*^*MiMIC*^ (**H**) and *Pvf3*^*MiMIC*^*/df*^*BSC291*^ (**I**) larvae compared to heterozygous controls (*hmlΔ-GAL4>UAS-GFP*, n≥7). (**J**) *Pvf2*^*T2A-GAL4*^*/+* heterozygotes have fewer blood cells than wildtype larvae (*hmlΔdsRed*, n=8). ns: not significant, lowercase letters indicate genotypes that differ significantly by an unpaired t-test with Welsh’s correction (**B-C**) or a Mann-Whitney test (**D-J**). Data points are individual larvae with means plotted ±1 standard error. Scale bars are 1mm.

Since macrophages represent >90% of the total population of blood cells, we next quantified total blood cells (circulating and tissue-resident) as a read out of macrophage number in *Pvf1-3* mutant larvae. As *Pvf1* is on the X chromosome we examined larvae hemizygous for the *Pvf1*^*1624*^ allele and found no defects in larval blood cell number (*hmlΔdsRed;* Figure 1D; p>0.999). In contrast, homozygous *Pvf2*^*T2A-GAL4*^ and transheterozygous *Pvf2*^*T2A-GAL4*^*/df*^*BSC291*^ larvae (where *df*^*BSC291*^ is a deficiency allele that deletes a number of genes including *Pvf2* and *Pvf3*, used to eliminate genetic background effects) both had less than one third the number of blood cells of the heterozygote controls (*hmlΔdsRed*; Figure 1E, F; for *Pvf2*^*T2A-GAL4*^, p<0.001 compared to *Pvf2*^*T2A-GAL4*^*/+*; for *Pvf2*^*T2A-GAL4*^*/df*^*BSC291*^, p<0.001 compared to both *Pvf2*^*T2A-GAL4*^*/+* and *df*^*BSC291*^*/+*). Similarly, homozygous *Pvf3*^*MiMIC*^ (*hml*Δ*-GAL4,UAS-GFP*), transheterozygous *Pvf3*^*MiMIC*^*/df*^*BSC291*^ (*hml*Δ*-GAL4,UAS-GFP*), and transheterozygous *Pvf3*^*MiMIC*^*/Pvf3*^*T2A-GAL4*^ (*hmlΔdsRed*) larvae all exhibited large reductions in blood cell number compared to their respective heterozygous controls (Figure 1G-I, S1; for *Pvf3*^*MiMIC*^, p<0.001 compared to *Pvf3*^*MiMIC*^*/+*; for *Pvf3*^*MiMIC*^*/df*^*BSC291*^, p<0.001 compared to both *Pvf3*^*MiMIC*^*/+* and *df*^*BSC291*^*/+*; for *Pvf3*^*MiMIC*^*/Pvf3*^*T2A-GAL4*^, p<0.001 compared to *Pvf3*^*T2A-GAL4*^*/+* and p=0.013 compared to *Pvf3*^*MiMIC*^*/Pvf3*^*T2A-GAL4*^). Together, these data indicate that *Pvf2* and *Pvf3*, but not *Pvf1*, influence larval blood cell number in a non-redundant manner. Given the magnitude of these reductions in blood cell number, it is likely that they predominantly reflect reductions to numbers of macrophages.

While quantifying *Pvf3*^*MiMIC*^*/df*^*BSC291*^ larvae, we noticed that *df*^*\BSC291*^*/+* control larvae had significantly fewer blood cells compared to *Pvf3*^*MiMIC*^*/+* controls (Figure 1I, p=0.019) and wondered whether this was because of haploinsufficiency for *Pvf2*. In support of this idea, we found that *Pvf2*^*T2A-GAL4*^*/+* individuals, which have ∼50% reduction in *Pvf2* transcript (Figure 1B), also have significantly reduced blood cell numbers compared to the wildtype control (Figure 1J, p=0.013). Unlike *Pvf2*^*T2A-GAL4*^ heterozygotes, *Pvf3*^*MiMIC*^*/+* larvae had a similar number of blood cells compared to wildtype controls (Figure 1H, p=0.336), despite similarly having a reduction in *Pvf3* transcript levels compared to wildtype animals (Figure 1C). This indicates that larval macrophage numbers may be more sensitive to dosage of *Pvf2* than *Pvf3*.

We also surveyed the crystal cell population to see if it was affected. We found crystal cell numbers to be strongly reduced in *Pvf2*^*T2A-GAL4*^ and *Pvf3*^*MiMIC*^ homozygotes, as well as in transheterozygotes of these alleles with the *df*^*BSC291*^ allele (Figure S2A-E). These data strongly implicate *Pvf2* and *Pvf3* in the control of crystal cell numbers. Since crystal cells transdifferentiate directly from macrophages during the larval stage (*23*), this is the likely the result of a compromised macrophage population in *Pvf2* and *Pvf3* mutants, rather than a separate role in crystal cells.

### *Pvf2* is required for larval macrophage self-renewal

Pvr is required for the survival of embryonic blood cells and has been proposed to be activated by all Pvfs in this role (*28, 42, 43*), however their relative contributions have not been resolved. The decreased number of macrophages in *Pvf2* and *Pvf3* mutant larvae could therefore be due to defects in the hematopoiesis that occurs in the embryo to establish the initial macrophage population in early larvae. Alternatively, the reduction could be due to defects in the self-renewal that occurs later in larval macrophages, or to reductions in both.

To determine which of these possibilities is occurring we therefore assessed blood cell numbers in first instar larvae (*hmlΔdsRed*), prior to the major macrophage self-renewal period and thus where blood cell numbers are determined by embryonic hematopoiesis. *Pvf2*^*T2A-GAL4*^*/Pvf2*^*T2A-GAL4*^ larvae exhibited no defects in blood cell number at this stage (Figure 2A, p=0.328). In contrast, *Pvf3*^*MiMIC*^*/Pvf3*^*T2A-GAL4*^ larvae had approximately half the number of blood cells compared to their heterozygous controls (Figure 2B, compared to *Pvf3*^*MiMIC*^*/+*: p<0.001; to *Pvf3*^*T2A-GAL4*^*/+*: p=0.005). When we estimated blood cell expansion rates using these data together with data from late stage larvae, we found no difference between *Pvf3*^*MiMIC*^*/Pvf3*^*T2A-GAL4*^ larvae and heterozygous controls (Figure S3A; Table S1), indicating that the phenotype observed in *Pvf3* mutants is solely due to an earlier role. Together, these data strongly suggest that Pvr signalling via Pvf2 but not Pvf3 mediates macrophage population expansion during the larval period.

**Figure 2.**
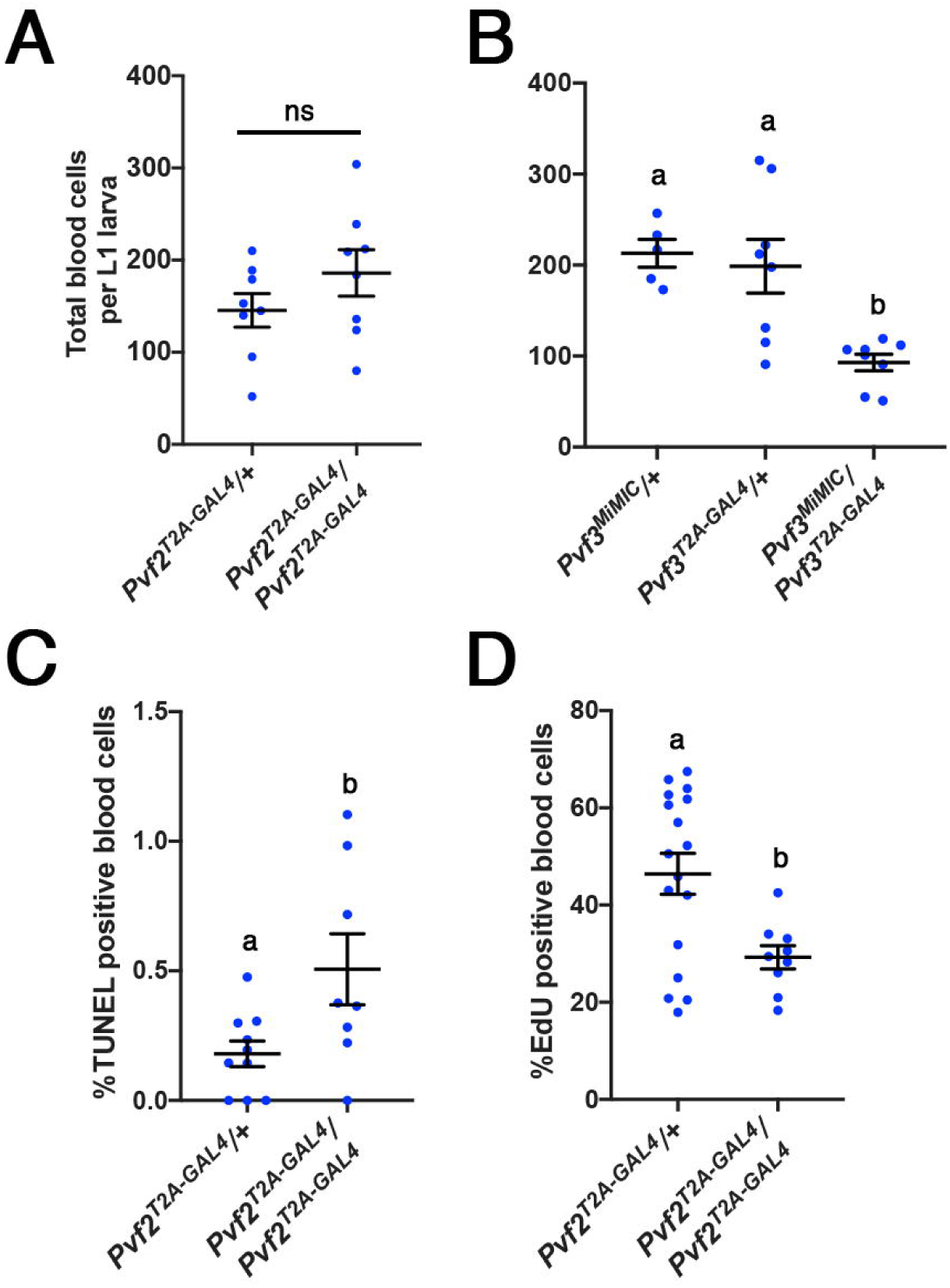
*Pvf2* mediates larval macrophage self-renewal. Total blood cell numbers in *Pvf2*^*T2A-GAL4*^*/Pvf2*^*T2A-GAL4*^ (**A**) and *Pvf3*^*MiMIC*^*/ Pvf3*^*T2A-GAL4*^ (**B**) larvae at the first instar stage. While *Pvf2*^*T2A-GAL4*^*/Pvf2*^*T2A-GAL4*^ larvae were not different from the heterozygous controls (n=8), *Pvf3*^*MiMIC*^*/ Pvf3*^*T2A-GAL4*^ larvae had markedly fewer blood cells (*hmlΔdsRed*, n≥5). (**C**) The percentage of TUNEL-positive blood cells is higher in *Pvf2*^*T2A-GAL4*^*/Pvf2*^*T2A-GAL4*^ larvae compared to heterozygous controls (n≥8). (**D**) Rates of blood cell DNA replication (EdU incorporation) are strongly reduced in *Pvf2*^*T2A-GAL4*^*/Pvf2*^*T2A-GAL4*^ larvae compared to heterozygous controls (n≥9). ns: not significant, lowercase letters indicate genotypes that differ significantly by a Mann-Whitney test. Data points are individual larvae with means plotted ±1 standard error.

Due to our interest in macrophage self-renewal, we focussed our attention on investigating how *Pvf2* controls the larval macrophage population. We reasoned that *Pvf2* may achieve this by promoting macrophage survival and/or self-renewal. To assess survival, we measured apoptosis in *Pvf2* mutants early during the third instar stage when cell numbers are rapidly increasing. While the overall occurrence of blood cell death was very low at this stage in both *Pvf2*^*T2A-GAL4*^ homozygotes and heterozygous controls (0-1.2%), the effect was significantly greater upon loss of *Pvf2* (Figure 2C; p=0.045). However, such low levels of cell death were unlikely to explain the severe reduction in macrophages that we observed in *Pvf2* mutants, so we suspected that there was a self-renewal deficit. We therefore assessed the rate of cell cycle progression by determining the rate at which blood cells incorporate the thymidine analogue EdU. Blood cells in *Pvf2*^*T2A-GAL4*^ homozygotes exhibited a marked and significant reduction in self-renewal rates (37%; Figure 2D; p=0.029). This aligns closely with the reduced blood cell expansion rate calculated by combining data from early and late stage larvae (35%, Figure S3B). We therefore conclude that *Pvf2* predominantly drives the expansion of the larval macrophage population by promoting their self-renewal.

Macrophage population expansion can be affected by both the accumulation of blood cells in hematopoietic pockets and by overall organismal growth (*13, 38, 44*). We therefore tested if the reduction in macrophage number in *Pvf2* mutants could be due to defects in one of these processes. Defects in blood cell accumulation alter the relative proportion of blood cells in circulation (*25*), so we assessed this proportion in *Pvf2*^*T2A-GAL4*^*/df*^*BSC291*^ transheterozygotes. We found that the relative proportions of tissue-resident to circulating blood cells did not differ from heterozygous controls (Figure S4A). In addition, *Pvf2* mutant animals were not growth-impaired (Figure S4B) and their ability to coordinate blood cell numbers with body size was also not affected (Figure S4C; Table S2). These data suggest that *Pvf2* may regulate the self-renewal process via a novel mechanism.

### A subpopulation of blood cells express *Pvf2* for macrophage self-renewal

To better understand how *Pvf2* influences macrophage self-renewal, we next investigated the tissue/ cellular origin(s) of *Pvf2* for this role. For this we used the *Pvf2*^*T2A-GAL4*^ allele to drive expression of a *YFP* reporter and explored whether *Pvf2* is expressed in blood cells. We first validated that the *Pvf2*^*T2A-GAL4*^ reporter faithfully reproduced the reported expression patterns of *Pvf2* (Figure S5A, B). We then tested for expression in larval blood cell populations by determining whether YFP (*Pvf2*^*T2A-GAL4*^*/+>UAS-YFP*) colocalises with dsRed (*hmlΔdsRed*) in larvae. This revealed a YFP signal that was significantly above background levels in a small number of both tissue-resident and circulating blood cells from *Pvf2*^*T2A-GAL4*^*/+>UAS-YFP* larvae (∼1%, Figure 3A; for tissue-resident, p=0.001 compared to both *Pvf2*^*T2A-GAL4*^*/+* and *UAS-YFP*; for circulating, p=0.019 compared to *Pvf2*^*T2A-GAL4*^*/+* and p=0.010 compared to *UAS-YFP*). This was further confirmed by live-imaging, which showed that these cells, like other blood cells, are dispersed throughout the larva (Figure 3B, Movie 1). Thus, the Pvf2 required for larval macrophage self-renewal may be produced by the blood cells themselves.

**Figure 3.**
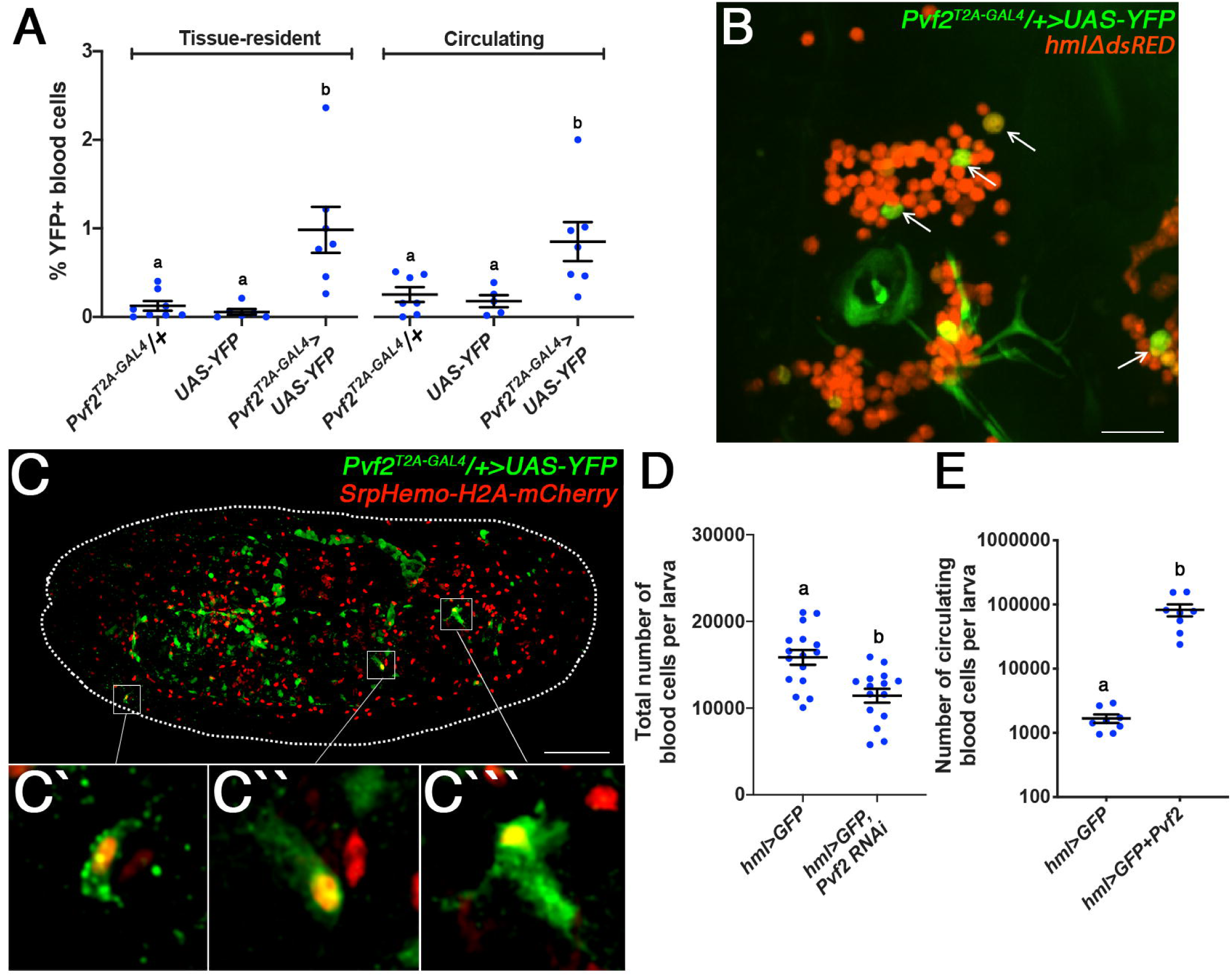
*Pvf2* expression in a blood cell subpopulation drives self-renewal. (**A**) Less than 1% of circulating and tissue-resident blood cells from *Pvf2*^*T2A-GAL4*^*/+>UAS-YFP* larvae express YFP above background levels (*Pvf2*^*T2A-GAL4*^/+ and *UAS*-*YFP* alone, n≥5). (**B**) Representative still image of the posterior-dorsal side of a live third instar *Pvf2*^*T2A-GAL4*^*/+>UAS-YFP* larva showing blood cells (*hmlΔdsRed*) including several that express YFP (arrowed). (**C**) Late-stage *Pvf2*^*T2A-GAL4*^*/+>UAS-YFP* embryo (anti*-*GFP to visualise YFP) showing a small number of blood cells (*SrpHemo-H2A-mCherry*, nuclear) that colocalise with the YFP signal. Magnifications of three YFP-positive blood cells are shown below (**C’**-**C’’’**). (**D**) Larvae expressing *Pvf2 RNAi* in all larval blood cells (*hml-GAL4>UAS-GFP*) exhibit a reduction in blood cell number (n≥15). (**E**) Overexpression of *Pvf2* in all larval blood cells (*hml-GAL4>UAS-GFP*) strikingly increases in circulating blood cell number (n=8). Lowercase letters indicate genotypes that differ significantly by a Mann-Whitney test. Data points are individual larvae with means plotted ±1 standard error. Scale bars are 50μm.

Notably, we did not detect *Pvf2* expression in blood cells within the lymph gland (Figure S5C), suggesting that *Pvf2* expression may be restricted to blood cells of embryonic origin. In agreement with this, when we looked at late-stage *Pvf2*^*T2A-GAL4*^*/+>UAS-YFP* embryos co-expressing an embryonic blood cell marker (*SrpHemo-H2A-mCherry*) (*45*), we consistently found several YFP-positive blood cells (∼1-5 per embryo; Figure 3C, Movie 2). This indicates that these cells are present throughout the larval stage. Consistent with *Pvf3* not playing a role in self-renewal, we did not detect *Pvf3* expression in larval blood cells (Figure S6A-C).

To determine if blood cell-produced Pvf2 is required for macrophage expansion, we knocked-down *Pvf2* expression specifically in larval blood cells via RNAi driven by *hml-GAL4*. This resulted in a significant reduction in larval blood cell number compared to the controls carrying *hml-GAL4* alone (p=0.001; Figure 3D, S7). These data suggest that *Pvf2* expression in blood cells is important for macrophage self-renewal. Moreover, because ectopic *Pvf2* expression in all larval blood cells results in a striking overexpansion phenotype (Figure 3E, p<0.001) (*44*), rates of macrophage self-renewal may be the product of not only *Pvf2* expression levels within blood cells, but also the number of blood cells that express *Pvf2*.

### Blood cell *Pvf2* expression is transient

We were next interested to understand whether the observed *Pvf2* expression defines a novel lineage of *Drosophila* blood cells or instead represents a transient state. To address this, we performed a lineage tracing experiment where we used *Pvf2*^*T2A-GAL4*^ to drive expression of a photoconvertible green fluorescent protein (*UAS-EOS-GFP*) that is cleaved upon UV irradiation, causing it to emit light in the red spectrum (EOS-GFP^RED^; Figure 4A) (*46*). *Pvf2*^*T2A-GAL4*^*/+>UAS-EOS-GFP* larvae were irradiated such that *Pvf2*-expressing macrophages and all their descendants are marked by EOS-GFP^RED^. Conversely, any cells that later switch on expression of *Pvf2* are marked by EOS-GFP alone (green). Live-imaging 24 hours after photoconversion revealed most *Pvf2-*expressing blood cells, but not other *Pvf2*-positive tissues (e.g. trachea), to express EOS-GFP alone (Figure 4B). These data suggest that blood cells can switch on *Pvf2* expression throughout the larval stage.

**Figure 4.**
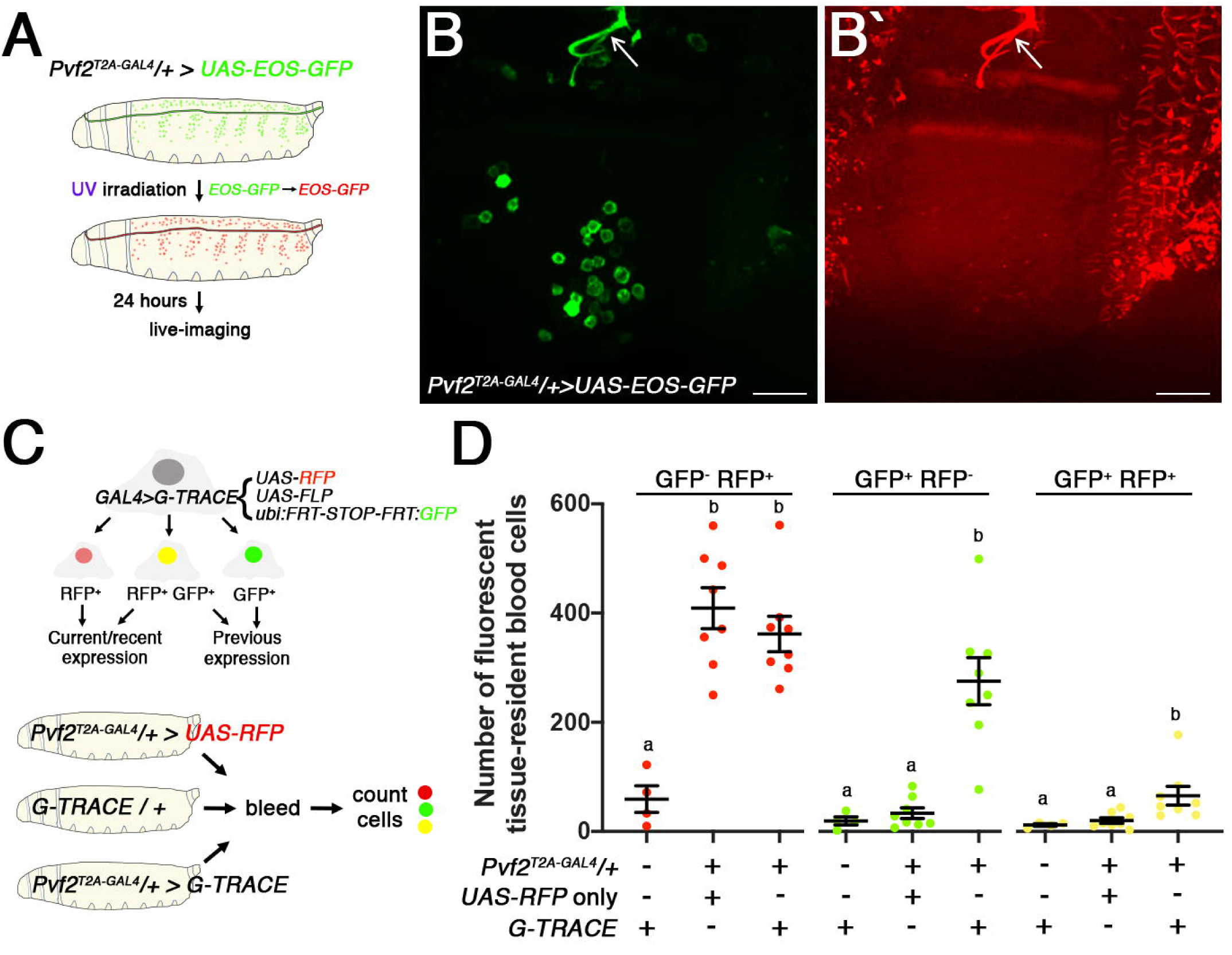
*Pvf2* expression in larval blood cells is a transient event. (**A**) Schematic of the EOS-GFP experiment. EOS-GFP (**B**, green channel) and UV-converted EOS-GFP (**B’**, EOS-GFP^RED^, red channel, overexposed) on the posterior-dorsal side of a live *Pvf2*^*T2A-GAL4*^*/+>UAS-EOS-GFP* larva 24 hours post-UV irradiation. Note that the blood cells contain EOS-GFP only, whereas the trachea (arrowed) also has EOS-GFP^RED^. (**C**) Schematic for the G-TRACE experiment. (**D**) Quantification of tissue-resident blood cells from *Pvf2*^*T2A-GAL4*^ /+, *Pvf2*^*T2A-GAL4*^*/+>RFP* and *Pvf2*^*T2A-GAL4*^*/+>G-TRACE* larvae. Larvae carrying both *Pvf2*^*T2A-GAL4*^*/+* and *G-TRACE* have RFP-and/ or GFP-positive blood cells that are above background levels (defined using controls not expressing the relevant fluorescent proteins; for RFP^+^GFP^-^, p=0.004 compared to larvae carrying *G-TRACE* alone; for RFP^-^GFP^+^, p=0.004 compared to larvae carrying *G-TRACE* alone and p<0.001 compared to *Pvf2*^*T2A-GAL4*^*/+>RFP* larvae; for RFP^+^GFP^+^, p=0.004 compared to larvae carrying *G-TRACE* alone and p=0.003 compared to *Pvf2*^*T2A-GAL4*^*/+>RFP* larvae). The presence of GFP^-^RFP^+^ and GFP^+^RFP^-^ blood cells from *Pvf2*^*T2A-GAL4*^*/+>G-TRACE* larvae indicates that the cells either currently express *Pvf2*, or previously expressed *Pvf2* and since have ceased, respectively. Lowercase letters indicate genotypes that differ significantly by a Mann-Whitney test for a given colour combination. Data points are individual larvae with means plotted ±1 standard error. Scale bars are 50μm.

As an alternative means of testing whether *Pvf2* expression is transient, we performed a different lineage tracing experiment using the G-TRACE system (comprising *UAS-RFP, UAS-FLP* and *Ubi*^*p63*^*-FRT-STOP-FRT-GFP* transgenes) (*47*) driven by *Pvf2*^*T2A-GAL4*^. Expression of FLP recombinase in *Pvf2*-expressing cells (marked by RFP) causes permanent expression of GFP in these cells and their descendants. Thus, here GFP is an indicator of previous expression and RFP an indicator of current or recent expression (Figure 4C). Extraction of the tissue-resident blood cells from *Pvf2*^*T2A-GAL4*^*/+>G-TRACE* larvae revealed the presence all combinations of RFP-and GFP-positive cells above background levels (Figure 4D, S8A, B). In particular, the presence of GFP-positive and RFP-negative blood cells in *Pvf2*^*T2A-GAL4*^*/+>G-TRACE* larvae strongly suggests that, during the larval stage, these cells cease expressing *Pvf2* (p=0.004 compared to larvae carrying *G-TRACE* only and p<0.001 compared to *Pvf2*^*T2A-GAL4*^*/+>RFP*). These data thus support the idea that *Pvf2*-expression in blood cells is a transient event.

## Discussion

Here we define a novel mechanism for the control of macrophage self-renewal in *Drosophila*, at the centre of which is the VEGF- and PDGF-related receptor (Pvr) signalling pathway. Pvr has been previously implicated in the control of blood cell number, however its contribution to their expansion in larvae and the nature of this have remained unknown (*28, 30, 31*). We found that Pvr is controlled by one of the three known Pvr ligands in this role, Pvf2. Specifically, we show that *Pvf2* expression levels in circulating and tissue-resident larval blood cells controls the rate of macrophage self-renewal and, to a lesser extent, blood cell survival. Our study however, does not rule out sources of Pvf2 in tissues other than the blood cells. Indeed, we noted its expression in various tissues that may supply Pvf2 to the lymph (e.g. trachea). However, we reason that the observed sensitivity of blood cells to more or less *Pvf2* argues strongly that the blood cells act as significant contributors to the pool of Pvf2 required for macrophage self-renewal.

Most notably our data suggests that, while only a small percentage of blood cells express *Pvf2*, these cells do not represent a novel blood cell lineage. Instead, this expression pattern is maintained via the transient expression of *Pvf2* in blood cells. Consistent with our findings, single cell RNA-sequencing of larval blood cells has shown that only a small proportion of blood cells express *Pvf2* (∼4%) and these cells do not cluster with any of the 16 identified blood cell lineages (Figure S9) (*48*). Currently, it remains unknown what signal(s) controls *Pvf2* expression in blood cells. However, since *Pvf2* expression is observed in blood cells from late embryogenesis and throughout the larval stage, the signal(s) that activates *Pvf2* expression is likely present during all of these stages. Whether this signal can influence *Pvf2*-expression in response to environmental stresses (e.g. infection, wounding) to modulate macrophage number is yet to be determined.

We have also shown that *Pvf3* determines the final macrophage population size. However, our data suggests that, unlike *Pvf2, Pvf3* is required prior to the onset of the larval stage and does not control larval macrophage self-renewal. In the embryo, Pvr influences blood cell numbers by promoting their survival, with genetic removal of Pvr resulting in up to 70% fewer blood cells (*28*). Although several studies have proposed that all Pvr ligands are required for the survival of embryonic blood cells, this has not been confirmed via the use of single *Pvf1-3* mutants (*28, 42, 43*). Our assessment of single mutants for *Pvf2* and *Pvf3* early during the larval stage revealed that *Pvf3* mutants exhibit a similar reduction in blood cell number to *Pvr* mutants (∼50%). In addition, the data reported here and the single cell RNA-sequencing conducted by Tattikotta et al. revealed that negligible numbers of larval blood cells express *Pvf3* (Figure S9) (*48*). Instead, *Pvf3* is more highly expressed in embryonic blood cells than larval blood cells (*49*). Together, these data suggest that *Pvf3* alone is likely responsible for Pvr-mediated blood cell survival in embryos.

Our data further suggests that the transition from embryonic to larval hematopoiesis involves a switch in Pvr function from supporting macrophage survival to promoting their self-renewal. Interestingly, this switch coincides with a change in the ligand that activates Pvr. What purpose the shifting of ligands serves to this and whether the promotion of larval macrophage survival has been relegated to another pathway (such as the Toll pathway) (*50*) remain interesting questions.

The data presented here describes and underscores the importance of the Pvr signalling pathway as a central regulator of *Drosophila* hematopoiesis. Importantly, our data reveal a new mechanism for how macrophage self-renewal can be regulated involving PDGF/VEGF receptor signalling. Members of these signalling pathways including the PDGFR-like colony stimulating factor receptor 1, and more recently VEGF-C, have been implicated in macrophage self-renewal in vertebrates (*4, 51, 52*). The strong similarities between *Drosophila* and vertebrates in this regard suggest that these cell signalling pathways likely have ancient evolutionary origins in the control of macrophage self-renewal. Indeed it will be interesting to learn the extent to which this is true and the degree to which regulatory mechanisms are conserved across distant animal species.

## Supporting information

Mov 1

Mov 2

## Acknowledgments

We thank the Australian *Drosophila* Biomedical Research Facility (OzDros) and Monash Micro Imaging for technical support. In addition, we thank the Bloomington *Drosophila* Stock Centre, Kyoto Stock Centre, and Katja Brückner for fly stocks, Linda Parsons, Tim Connallon, Christen Mirth and Grace Jefferies for advice and critical reading of the manuscript. J.C.W. is an Australian Research Council Laureate Fellow. This work was further supported by the Monash University Science-Medicine, Nursing, and Health Science Faculties Interdisciplinary Research Scheme.

## Movie 1

A third instar *Pvf2*^*T2A-GAL4*^*>UAS-YFP* larva with blood cells marked by *hmlΔdsRed*. Several YFP*-*expressing blood cells (yellow) are visible. Frames were captured at 30 second intervals for 58 minutes.

## Movie 2

A late-stage *Pvf2*^*T2A-GAL4*^*>UAS-YFP* embryo with blood cells marked by *SrpHemo-H2A-mCherry*. Note the YFP-positive blood cell indicated by an asterisk. Frames were captured at 2-minute intervals for one hour.

