## Supplementary material for "*Drosophila* macrophage self-renewal is regulated by transient expression of PDGF- and VEGF-related factor 2"

**Supplemental figures and legends**

**Figure S1.**

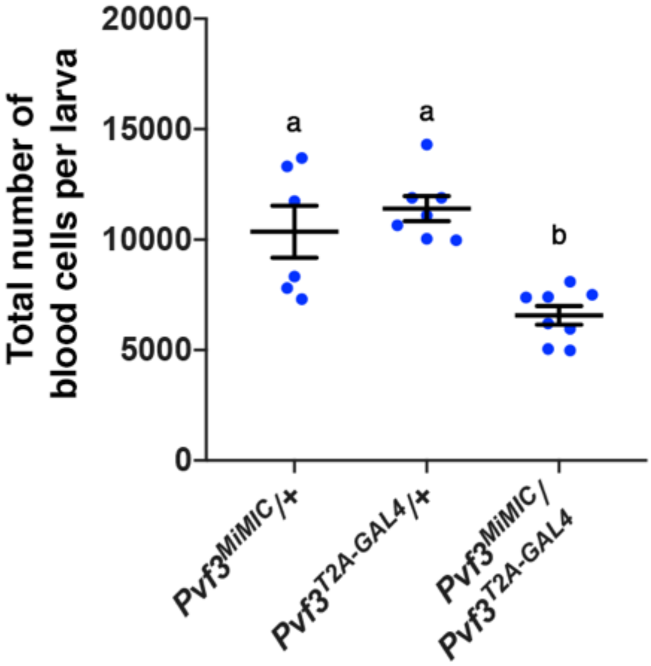


**Figure S1.** ***Pvf3*$^{T2A\text{-}GAL4}$ is a mutant allele of *Pvf3***

*Pvf3*$^{MiMIC}$/*Pvf3*$^{T2A\text{-}GAL4}$ larvae have fewer blood cells than both heterozygous controls, indicating that *Pvf3*$^{T2A\text{-}GAL4}$ is a hypomorphic allele of *Pvf3* (*hmlΔdsRed*, $n \geq 6$). Lowercase letters represent genotypes that differ significantly by a Mann-Whitney test. Data points are individual larvae with means plotted ±1 standard error.

**Figure S2.**

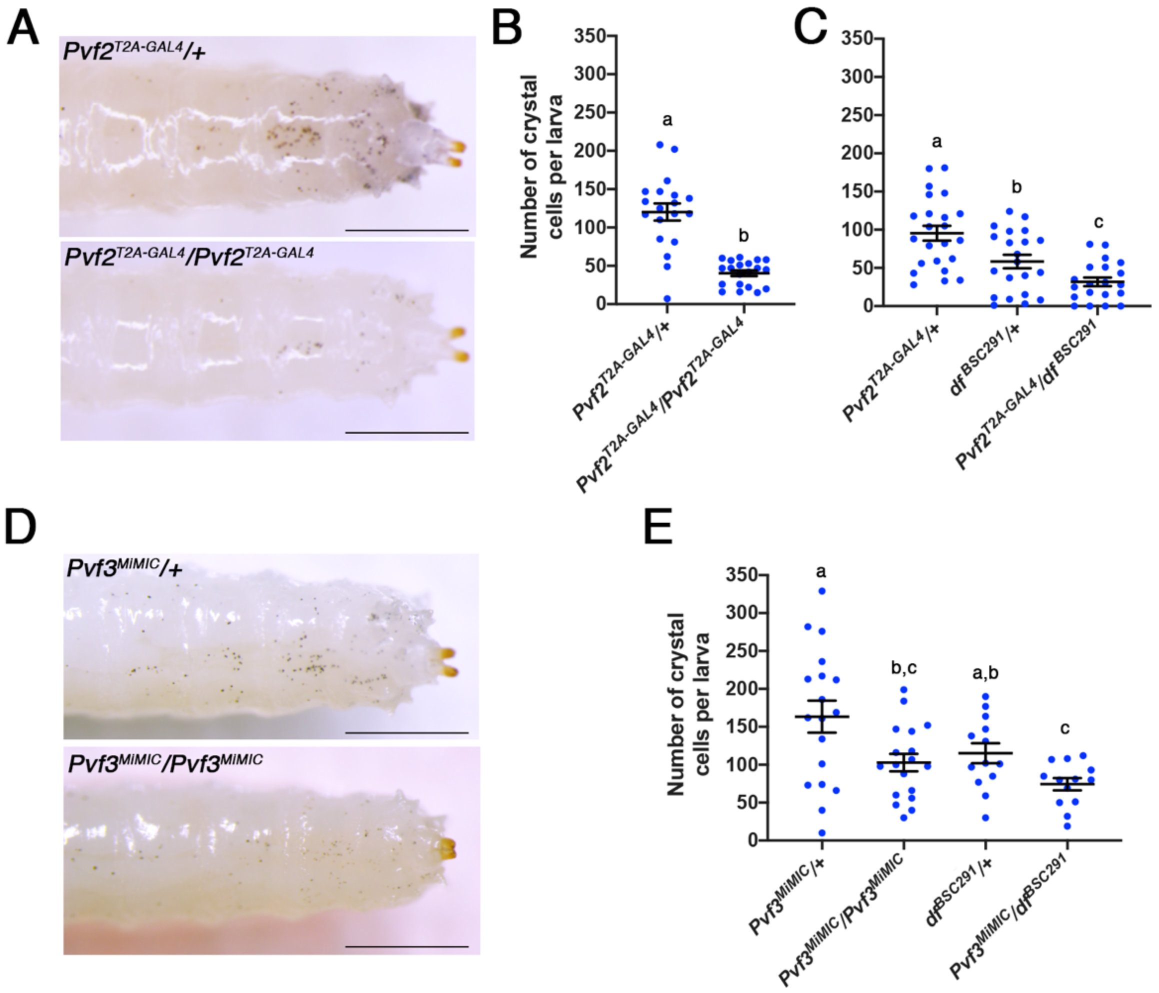


**Figure S2. *Pvf2* and *Pvf3* influence larval crystal cell numbers.**

(**A**) $Pvf2^{T2A\text{-}GAL4}/Pvf2^{T2A\text{-}GAL4}$ larvae appear to have fewer crystal cells (black puncta) compared to heterozygote controls. $Pvf2^{T2A\text{-}GAL4}/Pvf2^{T2A\text{-}GAL4}$ (**B**) and $Pvf2^{T2A\text{-}GAL4}/df^{BSC291}$ (**C**) have marked reductions in crystal cell numbers compared to heterozygous controls (for $Pvf2^{T2A\text{-}GAL4}/Pvf2^{T2A\text{-}GAL4}$, $p<0.001$ compared to $Pvf2^{T2A\text{-}GAL4}/+$; for $Pvf2^{T2A\text{-}GAL4}/df^{BSC291}$, $p<0.001$ compared to $Pvf2^{T2A\text{-}GAL4}/+$ and $p=0.029$ compared to $df^{BSC291}/+$, $n \geq 19$). (**D**) $Pvf3^{MiMIC}/Pvf3^{MiMIC}$ larvae appear to have fewer crystal cells compared to heterozygote controls. (**E**) $Pvf3^{MiMIC}/Pvf3^{MiMIC}$ and $Pvf3^{MiMIC}/df^{BSC291}$ larvae have fewer crystal cells than heterozygous controls (for $Pvf3^{MiMIC}/Pvf3^{MiMIC}$, $p=0.024$ compared to $Pvf3^{MiMIC}/+$; for $Pvf3^{MiMIC}/df^{BSC291}$, $p=0.008$ compared to $Pvf3^{MiMIC}/+$ and $p=0.031$ compared to $df^{BSC291}/+$,

$n \geq 13$). Lowercase letters represent genotypes that differ significantly by a Mann-Whitney test. Data points are individual larvae with means plotted $\pm 1$ standard error. Scale bars are 1mm.

**Figure S3.**

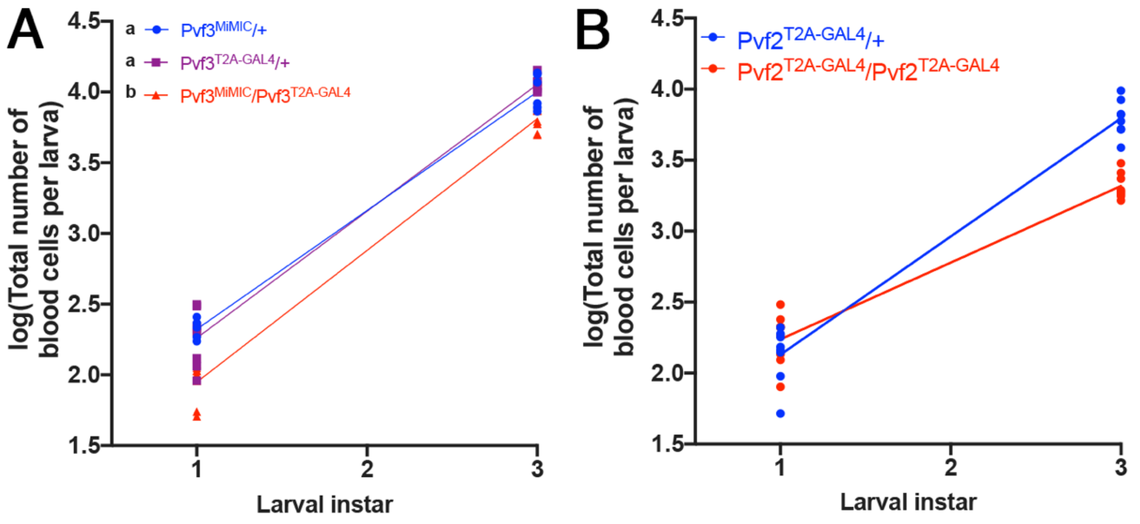


**Figure S3. Regression models of larval macrophage self-renewal by *Pvf2* and *Pvf3*.** Regression analysis of total blood cell number across larval stages in $Pvf3^{MiMIC}/Pvf3^{T2A\text{-}GAL4}$ transheterozygote (**A**) and $Pvf2^{T2A\text{-}GAL4}/Pvf2^{T2A\text{-}GAL4}$ (**B**) larvae compared to heterozygous controls (*hmlΔdsRed*, $n \geq 5$). (**A**) *Pvf3* mutant larvae have fewer blood cells than heterozygous controls at both stages ($p<0.001$ for both) such that the no difference was found in the blood cell expansion rate from first to third instars ($p=0.191$). Lowercase letters indicate significantly different regression lines. (**B**) $Pvf2^{T2A\text{-}GAL4}/Pvf2^{T2A\text{-}GAL4}$ larvae have fewer blood cells than heterozygous controls at the third instar larval stage, but not at the first instar stage (see main text). Regression lines for $Pvf2^{T2A\text{-}GAL4}/+$ ($Y = 0.8325*X + 1.297$) and $Pvf2^{T2A\text{-}GAL4}/Pvf2^{T2A\text{-}GAL4}$ ($Y = 0.5401*X + 1.698$) are shown. Note that the rate of expansion for $Pvf2^{T2A\text{-}GAL4}/Pvf2^{T2A\text{-}GAL4}$ is 35% less than the control, which aligns closely with the observed reduction in the rate of self-renewal in these larvae (37%, Figure 2D).

**Figure S4.**

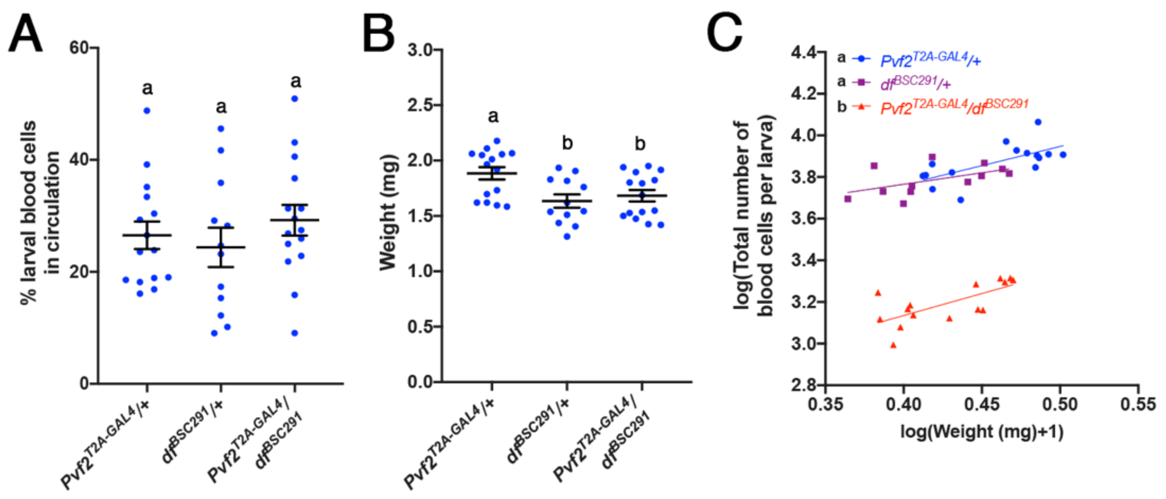


**Figure S4. *Pvf2* does not influence blood cell accumulation in hematopoietic pockets nor larval body size.**

Percent larval blood cells in circulation (**A**) and whole larva weights (**B**) of wandering $Pvf2^{T2A\text{-}GAL4}/df^{BSC291}$ larvae and heterozygous controls. $Pvf2^{T2A\text{-}GAL4}/df^{BSC291}$ larva have an equivalent percentage of circulating blood cells (p=0.389 compared to $Pvf2^{T2A\text{-}GAL4}/+$ and p=0.277 compared to $df^{BSC291}/+$) and body weight (p=0.614 compared to $df^{BSC291}/+$) compared to heterozygous controls (*hmlΔdsRed*, n≥12). (**C**) Regression analysis of total blood cell number per larva compared to the data in panel (**B**). $Pvf2^{T2A\text{-}GAL4}/df^{BSC291}$ larvae have fewer blood cells at any given final weight compared to the heterozygous controls (p<0.001 for both) and the relationship between weight and blood cell number is unaffected (p=0.447). Lowercase letters represent genotypes that differ significantly by a Mann-Whitney test (**A**, **B**) or significantly different regression lines (**C**). Data points are individual larvae with means plotted ±1 standard error.

**Figure S5.**

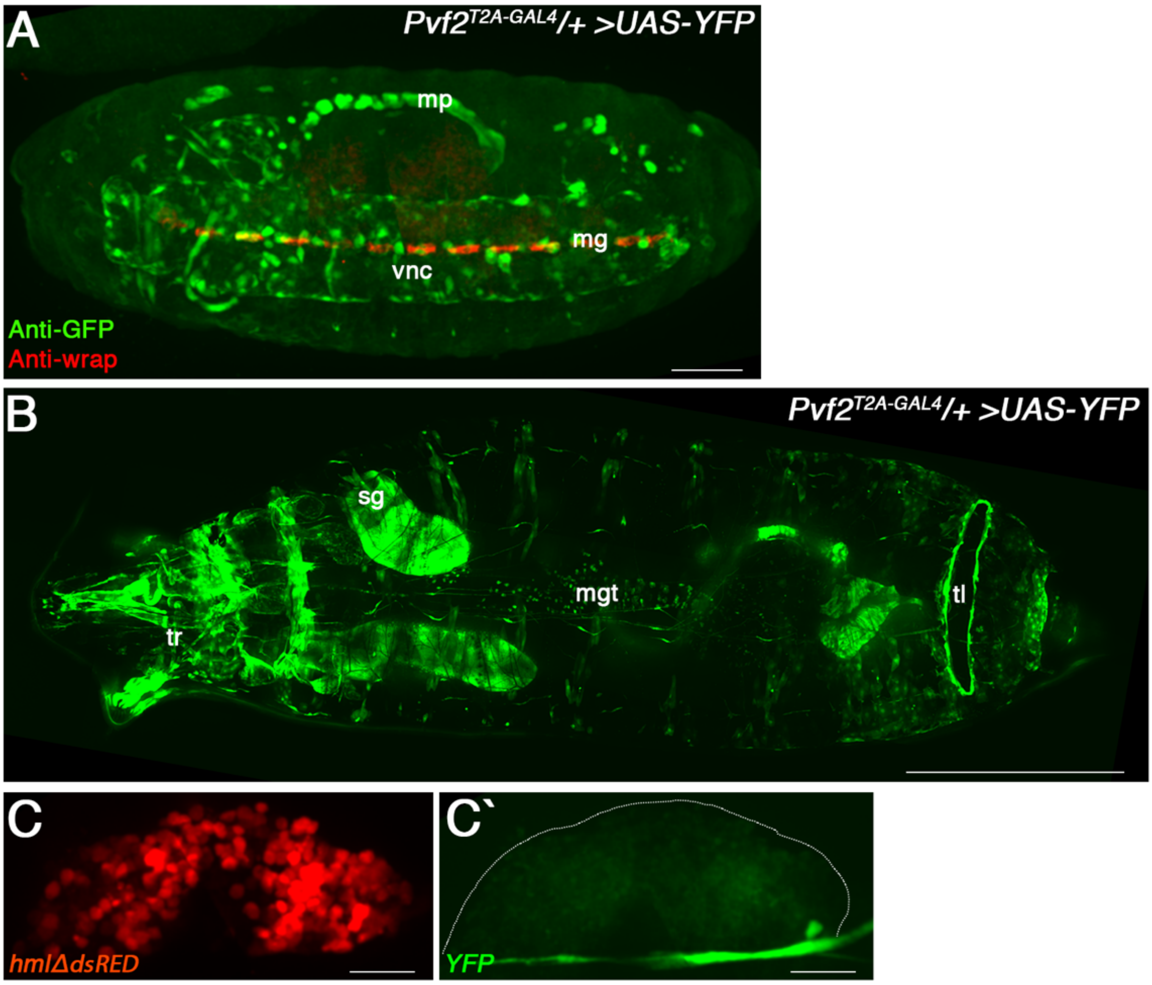


**Figure S5.** ***Pvf2**$^{T2A\text{-}GAL4}$* **expression in embryos and larvae.**

(**A**) Stage 16 embryo expressing YFP under the control of *Pvf2*$^{T2A\text{-}GAL4}$ immunostained with anti-GFP (green) and anti-wrapper (red). YFP expression is observed in the malpighian tubules (mp), ventral nerve cord (vnc) and midline glia (mg, anti-wrapper). This agrees with the previously observed pattern of *Pvf2* expression (*1*). Scale bar is 50µm. (**B**) Stitched maximum projection confocal image series of the ventral side of a live third instar larva. Notable expression is observed in the trachea (tr), salivary glands (sg), midgut (mgt) and telson (tl). This agrees with previously observed pattern of *Pvf2* expression (*2*). Scale bar is 100µm. (**C**) Representative lymph gland dissected from a wandering third instar *Pvf2*$^{T2A\text{-}GAL4}$*/+>UAS-eYFP* larva with blood cells marked by *hmlΔdsRed*. Note that cells of the aorta/heart express YFP, while blood cells do not. Scale bars are 50μm. Anterior is to the left.

**Figure S6.**

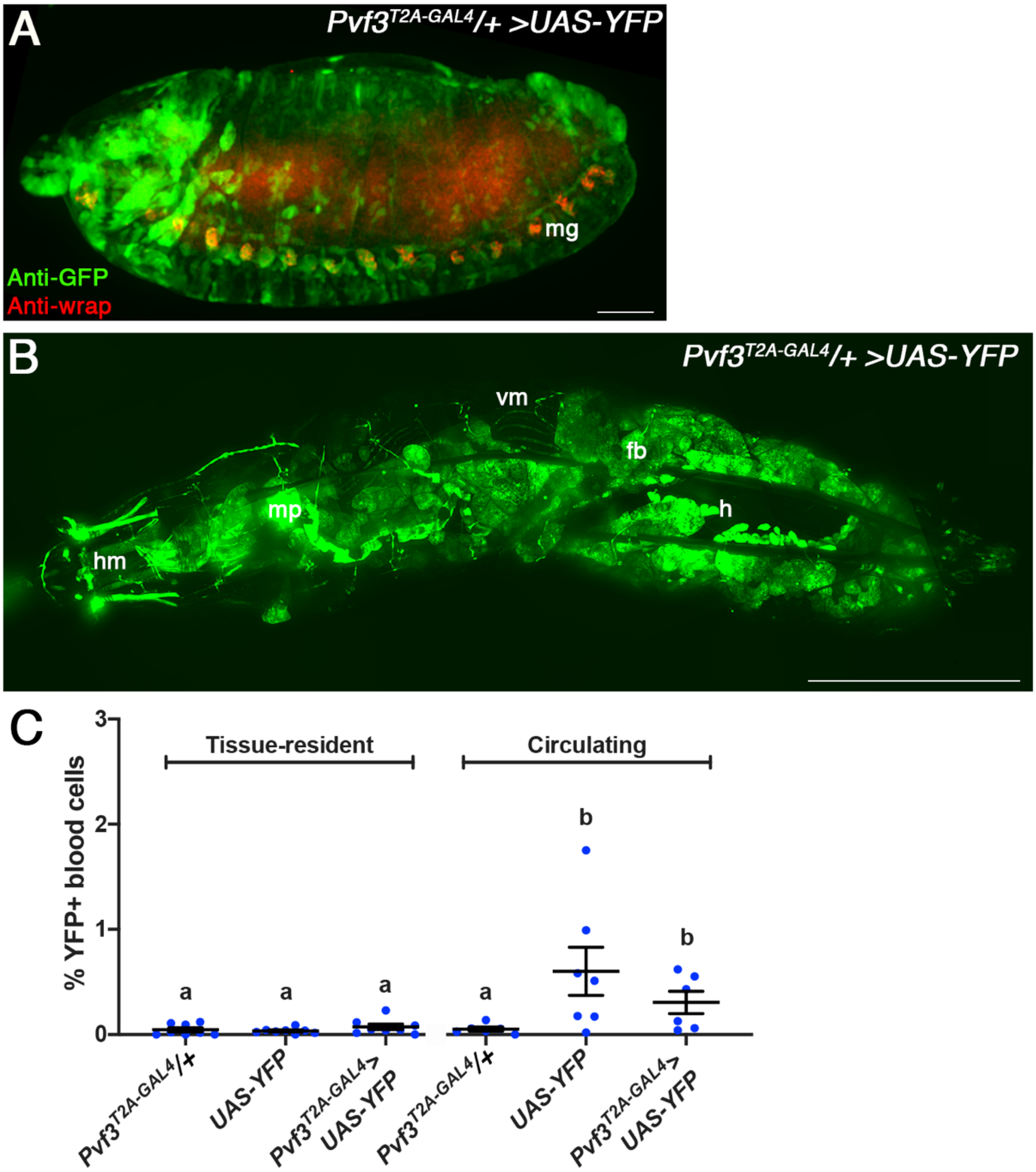


**Figure S6.** ***Pvf3**$^{T2A\text{-}GAL4}$* **expression in embryos and larvae.**

(**A**) Stage 14 embryo expressing YFP under the control of *Pvf3*$^{T2A\text{-}GAL4}$ immunostained with anti-GFP (green) and anti-wrapper (red). YFP expression is observed in midline cells of the ventral nerve cord including midline glia (mg, anti-wrapper), and extensively throughout the head. This agrees with previously observed pattern of *Pvf3* expression (*1*). Scale bar is 50µm. (**B**) Stitched maximum projection confocal image series of the dorsal side of a live third instar larvae. Notable expression is observed in the head musculature (hm), malpighian tubules (mp), vascular musculature (vm), fat body (fb) and heart (h). This agrees with previously observed pattern of *Pvf3* expression (*2*). Scale bar is 1mm. Anterior is to the left. (**C**) The percentage of YFP-positive circulating and tissue-resident blood cells detected in $\textit{Pvf3}^{\textit{T2A-GAL4}}$/+>*UAS-YFP* larvae does not significantly differ from background levels (controls not expressing *YFP*; for tissue-resident: p=0.454 compared to $\textit{Pvf3}^{\textit{T2A-GAL4}}$/+ and p=0.167 compared to *UAS-YFP*; for circulating: p=0.445 compared to *UAS-YFP*, n≥6). Lowercase letters indicate genotypes that differ significantly by a Mann-Whitney test. Data points are individual larvae with means plotted ±1 standard error.

**Figure S7.**

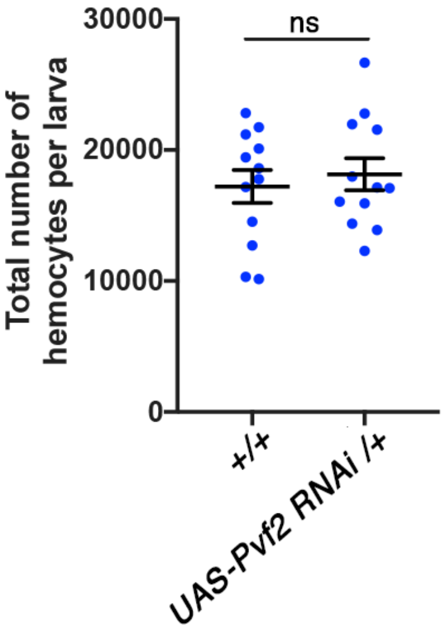


**Figure S7. Additional genetic background control for the *Pvf2 RNAi* experiment.**

The genetic background of *UAS-Pvf2 RNAi* does not influence blood cell numbers compared to the wildtype control (p=0.932, *hmlΔdsRed*, n=12). Ns: no significant difference, Mann-Whitney test. Data points are individual larvae with means plotted ±1 standard error.

**Figure S8.**

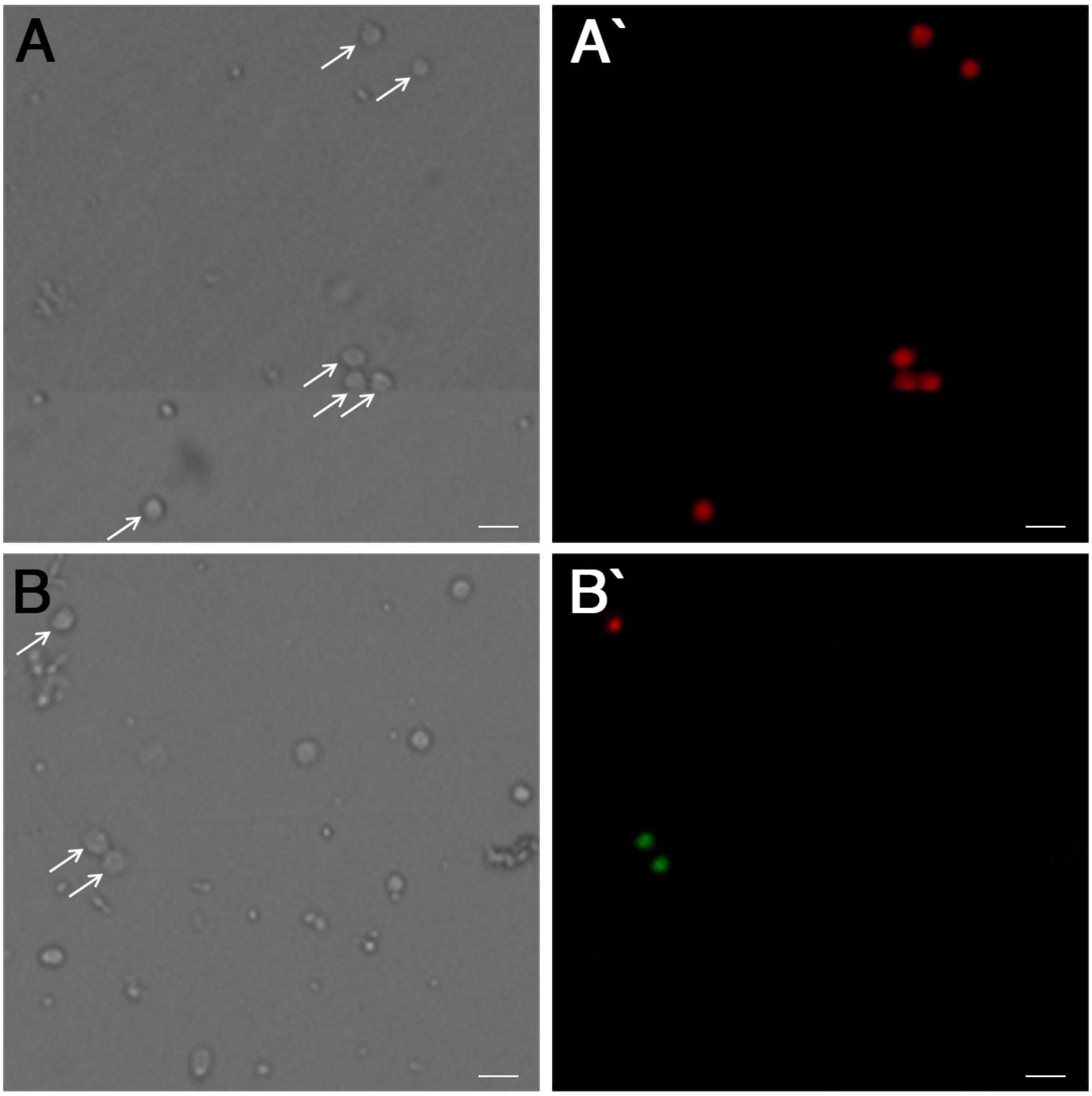


**Figure S8. G-TRACE positive cells extracted from *Pvf2*$^{T2A\text{-}GAL4}$/+>*G*-TRACE larvae are blood cells.**

Tissue-resident blood cells extracted from wandering *Pvf2*$^{T2A\text{-}GAL4}$/*hmlΔdsRed* (**A**, **A'**) and *Pvf2*$^{T2A\text{-}GAL4}$/+>*G-TRACE* (**B**, **B'**) larvae. Fluorescent cells are arrowed in brightfield images (**A**, **B**). The fluorescent signals detected in *Pvf2*$^{T2A\text{-}GAL4}$/+>*G-TRACE* larvae colocalise with blood cells rather than others cell types. Scale bars are 20μm.

**Figure S9.**

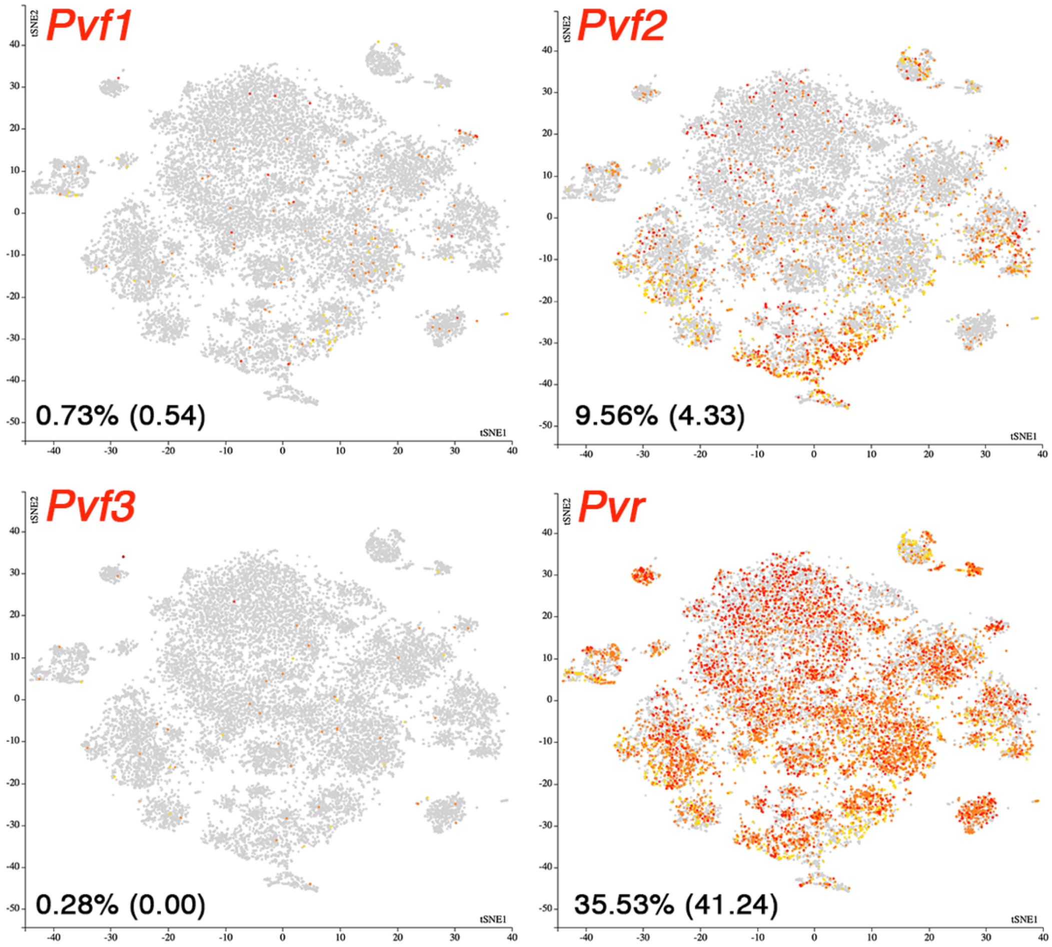


**Figure S9. Larval single blood cell RNA sequencing outputs for *Pvf1-3* and *Pvr* from Tattikota et al. (*3*).**

*Pvf1-3* and *Pvr* tSNE maps from single blood cell RNA sequencing analysis. Percentages of cells expressing each gene is indicated with unwounded values in parentheses. Maps are from integrated data combining unwounded, wounded and wasp-infected treatments courtesy of Tattikota et al. (https://www.flyrnai.org/scRNA/blood/) (*3*).

### Supplemental tables

**Table S1. Regression analysis statistics for data in Figure S3A.**

***$Pvf3^{T2A\text{-}GAL4}$/+* vs *$Pvf3^{MiMIC}$/+* vs *$Pvf3^{T2A\text{-}GAL4}$/ $Pvf3^{MiMIC}$***

| | Sum of squares | F value | Pr(>F) |
|---|---|---|---|
| **Stage** | 177.537 | 2095.763 | <0.001 *** |
| **Genotype** | 4.034 | 23.813 | <0.001 *** |
| **Stage*Genotype** | 0.0293 | 1.732 | 0.191 NS |

**Genotype Tukey post-hoc**

| | | T ratio | P value |
|---|---|---|---|
| ***$Pvf3^{T2A\text{-}GAL4}$/+* vs *$Pvf3^{MiMIC}$/+*** | | 0.094 | 0.995 NS |
| ***$Pvf3^{T2A\text{-}GAL4}$/+* vs *$Pvf3^{T2A\text{-}GAL4}$/ $Pvf3^{MiMIC}$*** | | -6.099 | <0.001 *** |
| ***$Pvf3^{MiMIC}$/+* vs *$Pvf3^{T2A\text{-}GAL4}$/ $Pvf3^{MiMIC}$*** | | 5.685 | <0.001 *** |

Data was fit with the linear model $\log_{10}$(total blood cell number) = Stage + Genotype + Stage*Genotype ($F_{5, 36}$ = 430.9, Adjusted $R^2$ = 0.981) using RStudio. This was followed with a Tukey posthoc test to determine which genotype(s) had significantly different effects on total blood cell number compared to the others.

**Table S2. Regression analysis statistics for data in Figure S4C.**

***$Pvf2^{T2A\text{-}GAL4}$/+* vs *$df^{BSC291}$/+* vs *$Pvf2^{T2A\text{-}GAL4}$/$df^{BSC291}$***

| | Sum of squares | F value | Pr(>F) |
|---|---|---|---|
| **$\log_{10}$(Weight)** | 0.660 | 25.825 | <0.001 *** |
| **Genotype** | 18.791 | 367.886 | <0.001 *** |
| **$\log_{10}$(Weight)*Genotype** | 0.042 | 0.823 | 0.447 NS |

**Genotype Tukey post-hoc**

| | | T ratio | P value |
|---|---|---|---|
| ***$Pvf2^{T2A\text{-}GAL4}$/+* vs *$df^{BSC291}$/+*** | | -0.820 | 0.693 NS |
| ***$Pvf2^{T2A\text{-}GAL4}$/+* vs *vs $Pvf2^{T2A\text{-}GAL4}$/$df^{BSC291}$*** | | 19.764 | <0.001 *** |
| ***$df^{BSC291}$/+ vs $Pvf2^{T2A\text{-}GAL4}$/$df^{BSC291}$*** | | 21.611 | <0.001 *** |

Data was fit with the linear model $\log_{10}$(total blood cell number) = $\log_{10}$(Weight) + Genotype + $\log_{10}$(Weight)*Genotype ($F_{5, 36}$ = 171.8, Adjusted $R^2$ = 0.954) using RStudio. This was followed with a Tukey posthoc test to determine which genotype(s) had significantly different effects on total blood cell number compared to the others.
